# *ATP7B* variant penetrance explains differences between genetic and clinical prevalence estimates for Wilson disease

**DOI:** 10.1101/499285

**Authors:** Daniel F. Wallace, James S. Dooley

**Author notes:** Corresponding author: (DFW).

## Abstract

Wilson disease (WD) is a genetic disorder of copper metabolism caused by variants in the copper transporting P-type ATPase gene *ATP7B*. Estimates for WD population prevalence vary with 1 in 30,000 generally quoted. However, some genetic studies have reported much higher prevalence rates. The aim of this study was to estimate the population prevalence of WD and the pathogenicity/penetrance of WD variants by determining the frequency of *ATP7B* variants in a genomic sequence database. A catalogue of WD-associated *ATP7B* variants was constructed and then frequency information for these was extracted from the gnomAD dataset. Pathogenicity of variants was assessed by (a) comparing gnomAD allele frequencies against the number of reports for variants in the WD literature and (b) using variant effect prediction algorithms. 231 WD-associated *ATP7B* variants were identified in the gnomAD dataset, giving an initial estimated population prevalence of around 1 in 2400. After exclusion of WD-associated *ATP7B* variants with predicted low penetrance, the revised estimate showed a prevalence of around 1 in 20,000, with higher rates in the Asian and Ashkenazi Jewish populations. Reanalysis of other recent genetic studies using our penetrance criteria also predicted lower population prevalences for WD in the UK and France than had been reported. Our results suggest that differences in variant penetrance can explain the discrepancy between reported epidemiological and genetic prevalences of WD. They also highlight the challenge in defining penetrance when assigning causality to some *ATP7B* variants.

## Introduction

Wilson disease (WD) is a rare autosomal recessive disorder of copper metabolism, resulting in copper accumulation with, most characteristically, hepatic and/or neurological disease (Ala et al. 2007). It is caused by variants in the gene encoding ATP7B, a copper transporter which in hepatocytes not only transports copper into the transGolgi for association with apocaeruloplasmin, but is fundamental for the excretion of copper into bile (Ala et al. 2007).

In WD, copper accumulates in the liver causing acute and/or chronic hepatitis and cirrhosis. Neuropsychiatric features are seen due to accumulation of copper in the brain. Other organs and tissues involved include the cornea (with the development of Kayser-Fleischer rings) and the kidneys.

There is a wide clinical phenotype and age of presentation. Early diagnosis and treatment are important for successful management (Ferenci et al. 2012). Diagnosis can be straightforward with a low serum ceruloplasmin associated with Kayser-Fleischer rings in the eyes, but may be difficult, requiring further laboratory tests, liver copper estimation and molecular genetic studies for *ATP7B* variants. Currently treatment of WD is either with chelators (d-penicillamine or trientine) which increase urinary copper excretion or zinc salts which reduce intestinal copper absorption (Ala et al. 2007; Ferenci et al. 2012). Liver transplantation may be needed for acute liver failure or decompensated liver disease unresponsive to treatment.

Diagnosis is often delayed (Merle et al. 2007) and there is a concern among clinicians that not all patients with WD are being recognised. It is not clear how great a problem this may be, but recent genetic studies have suggested a much higher prevalence of WD than is seen in population based epidemiological studies in the same country (Coffey et al. 2013; Collet et al. 2018; Park et al. 1991; Poujois et al. 2018). In addition a study using a global DNA dataset gave a much higher prevalence than generally accepted (Gao et al. 2019). Thus apart from data derived from some small isolated populations, where the reported incidence of WD is high (Dedoussis et al. 2005; Garcia-Villarreal et al. 2000), and small screening studies using low caeruloplasmin as the target (Hahn et al. 2002; Ohura et al. 1999), most epidemiological studies, including a study of all patients diagnosed in Denmark (Moller et al. 2011), predict prevalences that are in the range of 0.25 to 5.87 per 100,000 of the population (Gao et al. 2019). These figures are similar to the often quoted prevalence estimate of 30 per million from Scheinberg and Sternlieb in 1984 (Scheinberg and Sternlieb 1984).

Over 700 variants in *ATP7B* have been reported as associated with WD. The majority of patients are compound heterozygotes, the minority being homozygous for a single variant. A wide range of presentation and phenotype is recognised in WD and a relationship to *ATP7B* genotypes has been sought as a possible explanation, on the basis that causal variants will have different penetrance and expression. However, phenotype/genotype studies to date have shown a poor relationship (Chang and Hahn 2017; Ferenci et al. 2019). There has been increasing interest in the impact of other modifying genes and factors (Medici and Weiss 2017).

Thus questions remain regarding the biological impact of *ATP7B* variants on the clinical phenotype of WD and also its prevalence worldwide. The possibility of reduced penetrance of *ATP7B* variants has been suggested by recent studies (Loudianos et al. 2016; Sandahl et al. 2020; Stattermayer et al. 2019).

Next generation DNA sequencing (NGS) databases provide the opportunity to analyse the prevalence of WD variants in large populations and sub-populations. The Genome Aggregation Database (gnomAD) database contains variant frequencies derived from the whole exome or whole genome sequencing of over 120,000 people, from eight ethnic subgroups. NGS datasets are valuable resources and have been used by us and others for estimating the population prevalence of genetic diseases, such as HFE and non-HFE hemochromatosis (Wallace and Subramaniam 2016) and primary ubiquinone deficiency (Hughes et al. 2017). A recent study, that was published while this article was in preparation, used the gnomAD dataset to predict the prevalence of WD (Gao et al. 2019). The authors concluded that the global prevalence was around 1 in 7000 (Gao et al. 2019). We have taken a different approach to evaluate variant penetrance and predict that reduced penetrance of several *ATP7B* variants is likely to have a more substantial effect on the occurrence of clinically recognised WD, bringing the predicted prevalence from this NGS dataset closer to traditional estimates derived from epidemiologic studies (Scheinberg and Sternlieb 1984).

Our study highlights the difficulty in accurately assigning pathogenicity to *ATP7B* variants and the importance of defining penetrance when predicting the prevalence of inherited diseases using population genetic data. The resulting prevalence derived from this study is intermediate between historical estimates and those from more recent studies, at approximately 1 in 19,500, with figures above and below this in specific populations.

## Methods

### Catalogue of Wilson disease-associated *ATP7B* variants

Initially, details of all variants classified as “disease-causing variant” in the Wilson Disease Mutation Database (WDMD), hosted at the University of Alberta (http://www.wilsondisease.med.ualberta.ca/) were downloaded. As the WDMD has not been updated since 2010 a further literature search was carried out to identify WD-associated *ATP7B* variants that have been reported between the last update of the WDMD and April 2017, using the search terms ATP7B and mutation in the PubMed database (https://www.ncbi.nlm.nih.gov/pubmed). The Human Genome Variation Society (HGVS) nomenclature for each variant was verified using the Mutalyzer Name Checker program (https://mutalyzer.nl/) (Wildeman et al. 2008). Duplicate entries were removed and any mistakes in nomenclature were corrected after comparison with the original publications. All HGVS formatted variants were then converted into chromosomal coordinates (Homo sapiens – GRCh37 (hg19)) using the Mutalyzer Position Converter program. A variant call format (VCF) file containing all of the WD-associated *ATP7B* variants was then constructed using a combination of output from the Mutalyzer Position Converter and Galaxy bioinformatic tools (https://galaxyproject.org) (Afgan et al. 2016).

### Prevalence of Wilson disease-associated *ATP7B* variants

All variants in the *ATP7B* gene (Ensembl transcript ID ENST00000242839) were downloaded from the gnomAD (http://gnomad.broadinstitute.org/) browser (Lek et al. 2016). The WD-associated *ATP7B* variants (see above) were compared with the gnomAD *ATP7B* variants and allele frequency data were extracted for those variants with VCF descriptions that matched exactly. Allele frequency data were also extracted from the gnomAD dataset for variants that had not been previously reported in WD patients but were predicted to cause loss of function (LoF) of the ATP7B protein. These LoF variants included frameshift, splice acceptor, splice donor, start lost and stop gained variants.

Pathogenic *ATP7B* allele frequencies were determined in the gnomAD dataset by summing all of the allele frequencies for variants classified as WD-associated. Predicted pathogenic *ATP7B* genotype frequencies, heterozygote frequencies and carrier rates were calculated from allele frequencies using the Hardy-Weinberg equation.

### In silico analyses of variant pathogenicity

The functional consequence of WD-*ATP7B* missense variants and gnomAD-derived *ATP7B* missense variants (that had not been previously associated with WD) was assessed using the wANNOVAR program (http://wannovar.wglab.org/), which provides scores for 16 variant effect prediction (VEP) algorithms. The performance of these 16 algorithms for predicting WD-associated variants was analysed using receiver operating characteristic (ROC) curve analysis. The best performing algorithm (VEST3) (Carter et al. 2013) was used, together with the gnomAD frequency data, data from the WDMD and other published data (Gomes and Dedoussis 2016) to predict the pathogenicity of WD-associated *ATP7B* variants and refine the pathogenic genotype prevalence estimates.

## Results

### Wilson disease-associated *ATP7B* variants

The WDMD contained 525 unique *ATP7B* variants that have been reported in patients with WD and classified as disease causing (Supplementary Table S1). A literature search (between 2010 and April 2017) revealed a further 207 unique *ATP7B* variants associated with WD since the last update of the WDMD (Supplementary Table S2). Thus 732 *ATP7B* variants predicted to be causative of WD have been reported up until April 2017. For this study we refer to these 732 variants as WD-associated *ATP7B* (WD-*ATP7B*) variants.

The WD-*ATP7B* variants were categorized into their predicted functional effects, with the majority (400) being single base missense (non-synonymous) substitutions (Table 1). Variants predicted to cause major disruption to the protein coding sequence were further classified as loss of function (LoF). Variants were considered LoF if they were frameshift, stop gain (nonsense), start loss, splice donor, splice acceptor variants or large deletions involving whole exons. A total of 279 WD-*ATP7B* variants were categorized as LoF (Table 1) and their pathogenicity was considered to be high.

**Table 1.** Predicted functional consequences of WD-*ATP7B* variants.

| **Variant category** | **Number of variants** | **Loss of function (LoF)** | **Number in gnomAD** |
| --- | --- | --- | --- |
| Missense (non-synonymous) | 400 (55%) |  | 158 (68%) |
| Frameshift deletions, insertions or substitutions | 170 (23%) | Yes | 23 (10%) |
| Stop gain (nonsense) | 64 (9%) | Yes | 22 (10%) |
| Splice donor or acceptor | 43 (6%) | Yes | 10 (4%) |
| Non-frameshift deletions, insertions or substitutions | 26 (4%) |  | 4 (2%) |
| Intronic variants | 22 (3%) |  | 13 (6%) |
| Promoter variants | 2 (0.3%) |  |  |
| 5' UTR variants | 2 (0.3%) |  | 1 (0.4%) |
| Large deletions | 2 (0.3%) | Yes |  |
| Stop loss | 1 (0.1%) |  |  |
| Total | 732 |  | 231 |

### WD-*ATP7B* variants identified in the gnomAD dataset

Of the 732 WD-*ATP7B* variants 231 were present in the gnomAD dataset derived from >120,000 individuals (Lek et al. 2016) (Table 1). There was a higher proportion of missense variants among the WD-*ATP7B* variants present in gnomAD compared to the total WD-*ATP7B* variants reported in the literature (68% compared to 55%; Fisher’s Exact test, p=0.0002). Consequently there were also fewer LoF variants among the WD-*ATP7B* variants present in gnomAD (24% compared to 38%; Fisher’s Exact test, p<0.0001). However, we also identified an additional 51 LoF variants present in the gnomAD dataset that had not been reported in the literature as associated with WD (Supplementary Table S3). In addition we identified 10 copy number variants (CNVs), that would be expected to delete large portions of the *ATP7B* coding sequence, in the Exome Aggregation Consortium (ExAC) database, a forerunner of gnomAD, that contains approximately half the number of genomic sequences (Supplementary Table 3) (Ruderfer et al. 2016). We assumed that *ATP7B* LoF variants and CNV deletions would almost certainly be causative of WD when in the homozygous state or compound heterozygous state with other pathogenic *ATP7B* variants and their frequency data were included in subsequent calculations.

**Table 2.** Combined WD-*ATP7B* plus LoF variant allele frequencies, genotype frequencies and carrier rates in the gnomAD population.

|  | gnomAD |  |  |  |  |  |  |  |  |
| --- | --- | --- | --- | --- | --- | --- | --- | --- | --- |
|  | All | African | Ashkenazi Jewish | East Asian | European (non-Finnish) | European (Finnish) | Latino | South Asian | Other |
| Pathogenic allele freq | 0.02055 | 0.01245 | 0.03005 | 0.02369 | 0.02335 | 0.01775 | 0.01664 | 0.01591 | 0.02298 |
| Pathogenic genotype freq | 0.00042 | 0.00016 | 0.00090 | 0.00056 | 0.00055 | 0.00031 | 0.00028 | 0.00025 | 0.00053 |
| Heterozygous genotype freq | 0.04026 | 0.02459 | 0.05830 | 0.04627 | 0.04562 | 0.03486 | 0.03273 | 0.03131 | 0.04491 |
| Pathogenic genotype rate<sup>1</sup> | 2367 | 6451 | 1107 | 1781 | 1833 | 3176 | 3610 | 3952 | 1893 |
| Heterozygous carrier rate<sup>1</sup> | 25 | 41 | 17 | 22 | 22 | 29 | 31 | 32 | 22 |

<sup>1</sup> Pathogenic genotype rate and heterozygous carrier rate are expressed as 1 in “n” of the population.

**Table 3:**
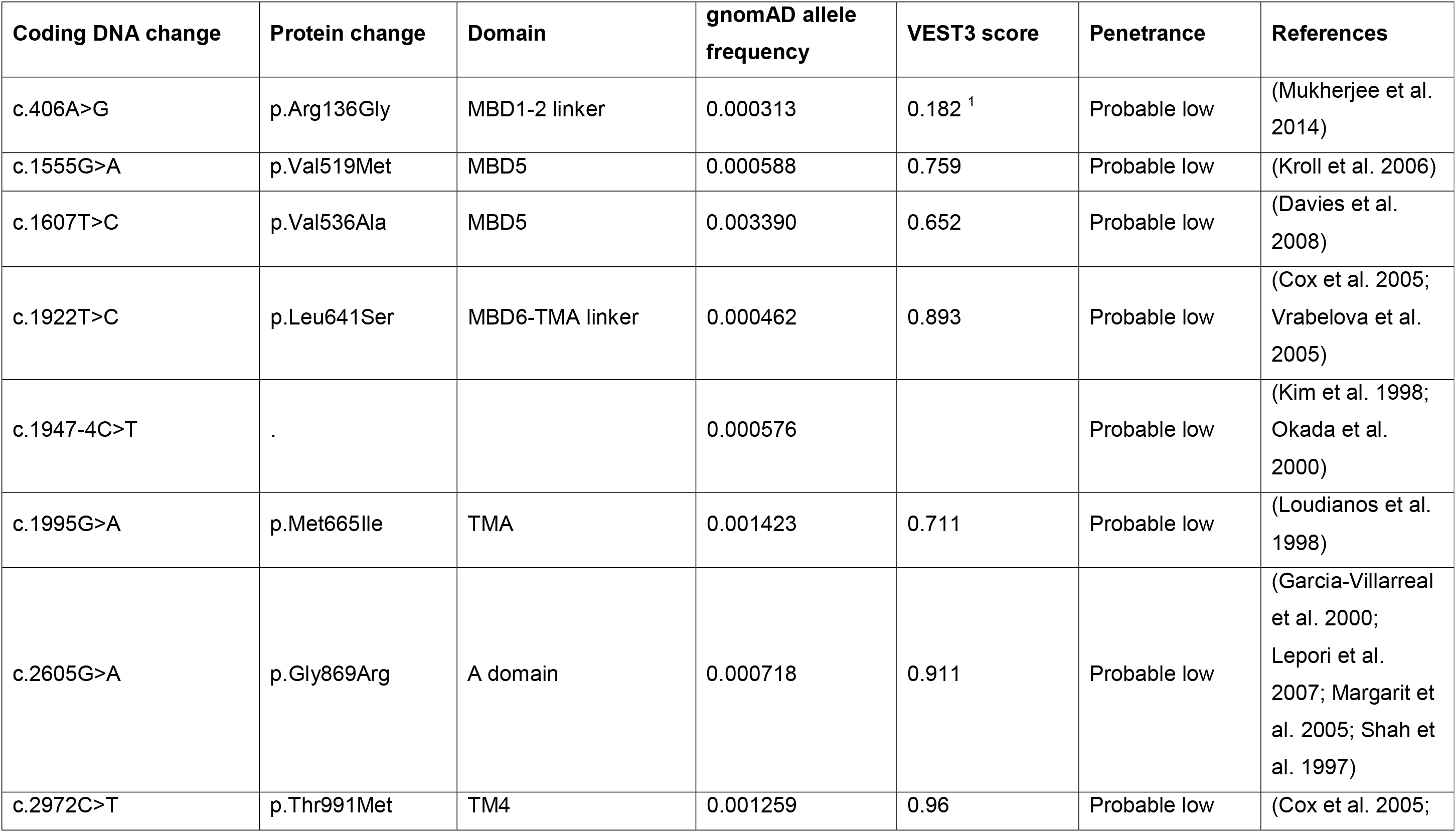
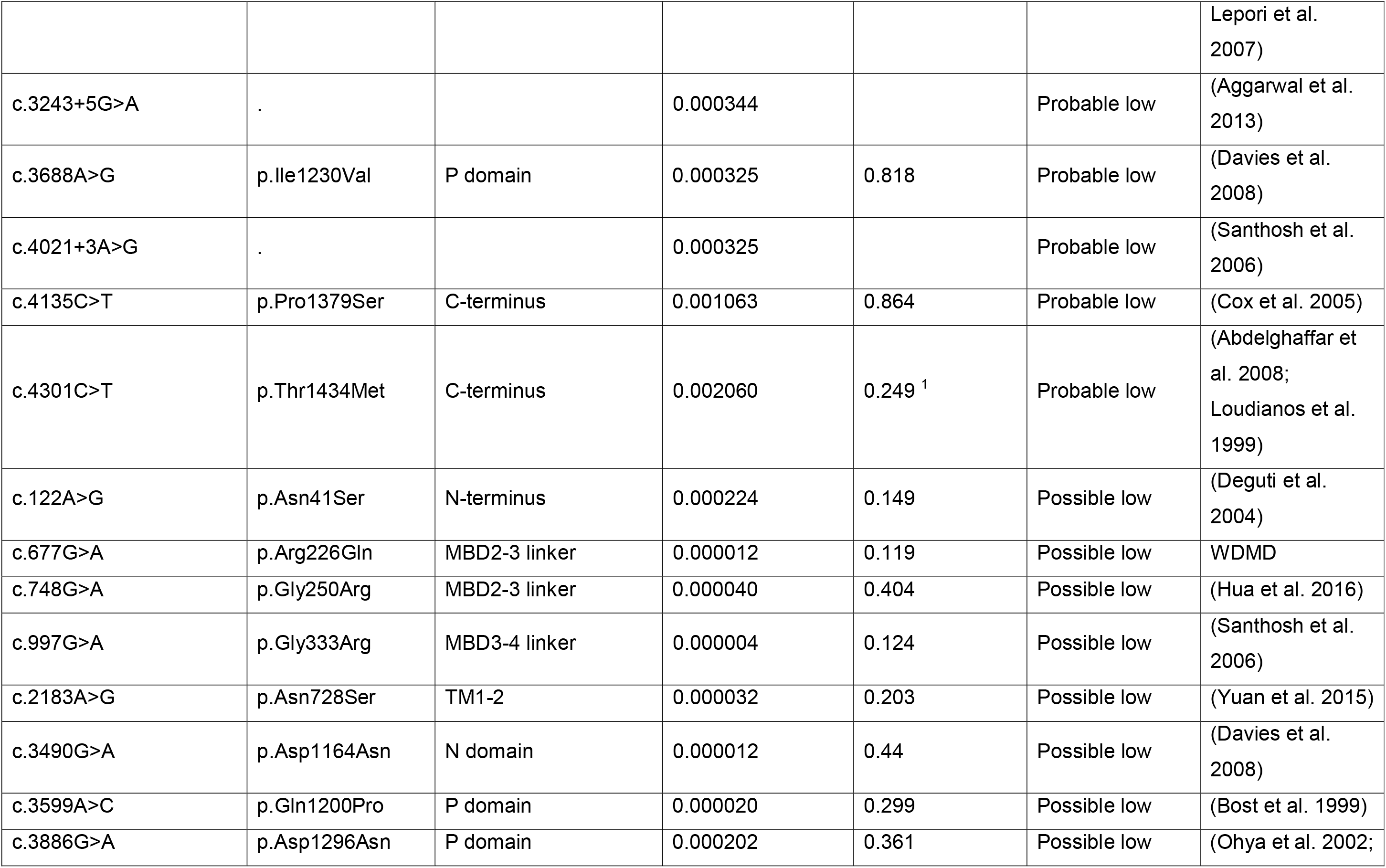
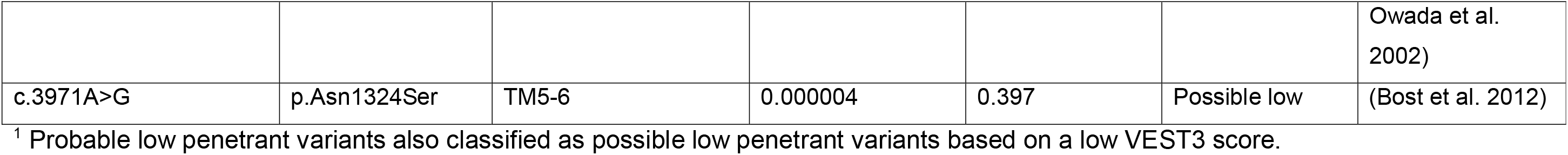
WD-*ATP7B* variants (found in the gnomAD dataset) with probable or possible low penetrance.

### Predicted prevalence of WD-associated genotypes in the gnomAD populations

Allele frequencies for all reported WD-*ATP7B* variants and the additional LoF variants and CNV deletions present in the gnomAD dataset were summed to give an estimate for the combined allele frequency of all WD-*ATP7B* variants in this population, which we have termed the pathogenic allele frequency (PAF). This was done for the entire gnomAD population and also for the 8 subpopulations that make up this dataset (Table 2). Assuming Hardy-Weinberg equilibrium and using the Hardy-Weinberg equation, the PAFs were used to calculate the pathogenic genotype frequencies (being homozygous or compound heterozygous for WD-*ATP7B* variants), the heterozygous genotype frequencies (being heterozygous for WD-*ATP7B* variants) and the carrier rates for these genotypes, expressed as one per “n” of the population (Table 2). The PAF in the whole gnomAD dataset was 2.055%, giving a pathogenic genotype rate (PGR) of 1 in 2367 and heterozygous carrier rate of 1 in 25. The highest PAF was seen in the Ashkenazi Jewish population (PAF 3.005%, PGR 1 in 1107) and the lowest in the African population (gnomAD: PAF 1.245%, PGR 1 in 6451). Frequency data were also calculated without the non-reported LoF variants and CNV deletions (Supplementary Table S4). However, due to the low combined allele frequency of these additional variants, the PAFs and PGRs were only marginally lower when these variants were not included (global population: PAF 2.004%, PGR 1 in 2491).

### Identification of low penetrant or non-causative *ATP7B* variants

Our initial estimate for the population prevalence of WD-*ATP7B* variants and consequently the predicted prevalence of WD in the gnomAD population of around 1 in 2400 with heterozygous carrier rate of 1 in 25 is considerably higher than the often quoted prevalence of 1 in 30,000 with 1 in 90 heterozygous carriers. It is also higher than prevalence estimates obtained from genetic studies in the UK (1 in 7000) (Coffey et al. 2013), France (1 in 4000) (Collet et al. 2018), and another recent study that also utilized allele frequency data from gnomAD (1 in 7000) (Gao et al. 2019). These genetic studies employed some filtering strategies, largely based on predictive software to remove *WD-ATP7B* variants that are likely to be benign or of uncertain significance. However, these genetic studies and our initial estimate do not appear to reflect the incidence of WD presenting to the clinic and suggest either that many WD patients remain undiagnosed or that some WD-*ATP7B* variants are not causative or have low penetrance.

We addressed the issue of variant penetrance using two approaches: firstly, by comparing the allele frequencies of individual variants in the gnomAD dataset with the frequency with which these variants have been reported in association with WD in the literature; and secondly by utilizing VEP algorithms.

In the first approach, if the allele frequency in the gnomAD dataset was such that more reports would have been expected in the literature (analysed broadly by number of references) then the variant was considered as a ‘*probable* low penetrant’ variant. Thus, when we ranked WD-*ATP7B* variants according to their allele frequencies in the gnomAD population we noticed that the p.His1069Gln variant, the most common WD-associated variant in European populations, was ranked number 6 in the entire gnomAD dataset, number 5 in the European (Finnish and non-Finnish) subpopulations and number 3 in the Ashkenazi Jewish subpopulation. Thus there were several WD-*ATP7B* variants with higher allele frequencies in these populations that would be expected to be detected regularly in WD patients. The 5 WD-*ATP7B* variants that ranked higher than p.His1069Gln in the gnomAD dataset were p.Val536Ala, p.Thr1434Met, p.Met665Ile, p.Thr991Met and p.Pro1379Ser. These variants have only been reported in a small number of cases of WD and hence their causality and/or penetrance is in question.

We also attempted to identify variants that have questionable causality/penetrance by comparing them against a recent review article that analysed the geographic distribution of *ATP7B* variants that have been reported in WD patients (Gomes and Dedoussis 2016). This review lists the most commonly encountered *ATP7B* variants in WD patients from geographic regions around the world. Any variants reported in this article were considered to have high penetrance. Interestingly, the 5 variants with gnomAD allele frequencies higher than p.His1069Gln were not listed in the Gomes and Dedoussis review (Gomes and Dedoussis 2016) suggesting that they are not commonly associated with WD.

We formalised this approach by analysing data from the WDMD. The WDMD lists all references that have reported particular variants. We counted the number of references associated with each WD-*ATP7B* variant (Supplementary Table S1). The p.His1069Gln variant is listed against 46 references, the highest number for any variant in the WDMD. In contrast the 5 variants with higher gnomAD allele frequencies have only 1 or 2 associated references in the WDMD (Supplementary Table S1), suggesting that their penetrance is low. We plotted gnomAD allele frequency against number of WDMD references for all WD-*ATP7B* variants and highlighted those variants that were reported by Gomes and Dedoussis (Gomes and Dedoussis 2016) (Figure 1A). This analysis showed that there were a number of variants with relatively high allele frequencies in gnomAD, not reported in the Gomes and Dedoussis review paper and with few references in the WDMD. These variants are clustered towards the left-hand side of the graph in Figure 1A. On the basis of this analysis we classified 13 variants as having ‘*probable* low penetrance’ (Table 3).

**Figure 1.**
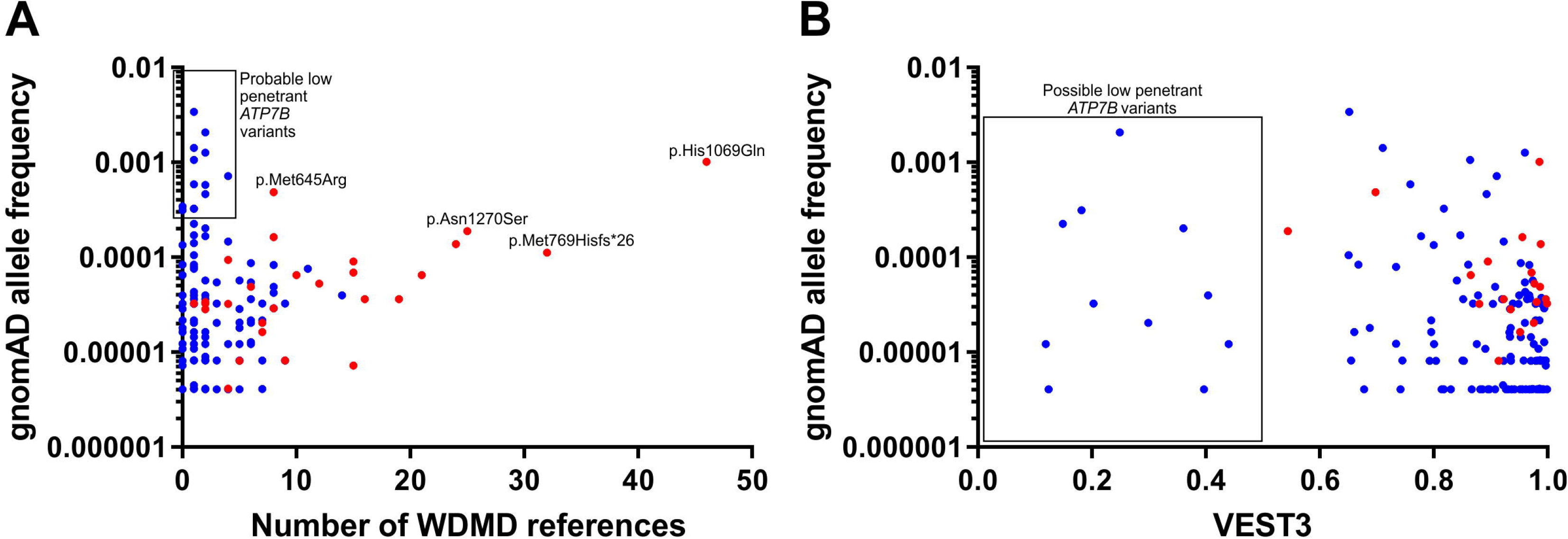
Identification of probable and possible low penetrant *ATP7B* variants. (A) The number of WDMD references was plotted against gnomAD allele frequency for WD-*ATP7B* variants identified in the gnomAD dataset. (B) VEST3 score was plotted against gnomAD allele frequency for WD-*ATP7B* variants identified in the gnomAD dataset. Variants reported in the Gomes and Dedoussis (Gomes and Dedoussis 2016) review as being the most common WD-*ATP7B* variants in various geographic regions are denoted by red dots and those not reported in the Gomes and Dedoussis review by blue dots. In (A) 13 variants were classified as *probable* low penetrant based on relatively high allele frequencies, low numbers of WDMD references and not being reported in the Gomes and Dedoussis review (boxed). In (B) 11 variants were classified as *possible* low penetrant based on a VEST3 score of <0.5 (boxed).

### Comparison of variant effect prediction (VEP) algorithms

VEP algorithms are used extensively to predict whether amino acid substitutions (missense variants) are likely to alter protein function and hence contribute to disease. We analysed the ability of 16 VEP algorithms to discriminate between WD-associated and non-WD-associated *ATP7B* missense variants (Supplementary Results). We determined that the VEST3 algorithm was the best at discriminating between WD and non-WD missense variants in the *ATP7B* gene (Supplementary Figures S1 and S2).

We classified WD-*ATP7B* missense variants found in the WDMD and in our literature search as ‘*possible* low penetrance’ if they had a VEST3 score of <0.5 (Figure 1B). There were 11 such variants in the gnomAD dataset that were contributing to our initial estimates of WD prevalence (Table 3). Two of these variants were also classified as *probable* low penetrance in the previous analysis based on the number of publications.

### Prevalence of WD-*ATP7B* variants in the gnomAD dataset after removing variants with probable or possible low penetrance

Exclusion from the analysis of the 13 WD-*ATP7B* variants with *probable* low penetrance, based on relatively high allele frequencies but low numbers of reports in WD patients, resulted in a significant reduction in the predicted prevalence of WD. The updated PAF after exclusion of these variants was 0.76% in the gnomAD dataset, with PGR of 1 in 16,832. The updated PAFs, genotype frequencies and carrier rates, including the results for each subpopulation can be seen in Table 4.

**Table 4.** Combined WD-*ATP7B* plus LoF variant allele frequencies, genotype frequencies and carrier rates in the gnomAD population after exclusion of those variants with *probable* low penetrance.

|  | gnomAD |  |  |  |  |  |  |  |  |
| --- | --- | --- | --- | --- | --- | --- | --- | --- | --- |
|  | All | African | Ashkenazi Jewish | East Asian | European (non-Finnish) | European (Finnish) | Latino | South Asian | Other |
| Pathogenic allele freq | 0.007708 | 0.003874 | 0.014093 | 0.015037 | 0.007884 | 0.004326 | 0.007724 | 0.006127 | 0.007902 |
| Pathogenic genotype freq | 0.000059 | 0.000015 | 0.000199 | 0.000226 | 0.000062 | 0.000019 | 0.000060 | 0.000038 | 0.000062 |
| Heterozygous genotype freq | 0.015297 | 0.007718 | 0.027790 | 0.029621 | 0.015645 | 0.008615 | 0.015328 | 0.012179 | 0.015680 |
| Pathogenic genotype carrier rate<sup>1</sup> | 16832 | 66625 | 5035 | 4423 | 16086 | 53427 | 16763 | 26636 | 16014 |
| Heterozygous carrier rate<sup>1</sup> | 65 | 130 | 36 | 34 | 64 | 116 | 65 | 82 | 64 |

<sup>1</sup> Pathogenic genotype rate and heterozygous carrier rate are expressed as 1 in “n” of the population.

The remaining 9 variants with *possible* low penetrance based on VEST3 score had lower allele frequencies and consequently their exclusion from the analyses had less effect on the predicted prevalence of WD. After exclusion of these variants the updated PAF decreased to 0.71% for the gnomAD dataset, with PGR of 1 in 19,457. The updated PAFs, genotype frequencies and carrier rates, including the results for each subpopulation can be seen in Table 5.

**Table 5.** Combined WD-*ATP7B* plus LoF variant allele frequencies, genotype frequencies and carrier rates in the gnomAD population after exclusion of those variants with *probable* and *possible* low penetrance.

|  | gnomAD |  |  |  |  |  |  |  |  |
| --- | --- | --- | --- | --- | --- | --- | --- | --- | --- |
|  | All | African | Ashkenazi Jewish | East Asian | European (non-Finnish) | European (Finnish) | Latino | South Asian | Other |
| Pathogenic allele freq | 0.007169 | 0.003583 | 0.014093 | 0.013720 | 0.007390 | 0.002613 | 0.007694 | 0.005932 | 0.007438 |
| Pathogenic genotype freq | 0.000051 | 0.000013 | 0.000199 | 0.000188 | 0.000055 | 0.000007 | 0.000059 | 0.000035 | 0.000055 |
| Heterozygous genotype freq | 0.014235 | 0.007140 | 0.027790 | 0.027064 | 0.014672 | 0.005212 | 0.015269 | 0.011794 | 0.014765 |
| Pathogenic genotype rate<sup>1</sup> | 19457 | 77903 | 5035 | 5312 | 18309 | 146467 | 16893 | 28415 | 18076 |
| Heterozygous carrier rate<sup>1</sup> | 70 | 140 | 36 | 37 | 68 | 192 | 65 | 85 | 68 |

<sup>1</sup> Pathogenic genotype rate and heterozygous carrier rate are expressed as 1 in “n” of the population.

## Discussion

We have used publically available NGS data firstly to predict the genetic prevalence of WD and secondly to assess the penetrance of WD variants to derive a more realistic WD prevalence that takes into account low penetrant variants.

Our initial estimates for population prevalence of WD included frequencies of all variants that had been reported as disease causing in the WDMD and more recent literature, with no adjustments for penetrance. We defined variants as WD-*ATP7B* variants if they were classified as disease causing in the WDMD or were reported as disease causing in the literature. Many of these variants could be defined according to the American College of Medical Genetics and Genomics (ACMG) guidelines (Richards et al. 2015) as pathogenic or likely pathogenic, however, for some there would be a lack of supporting information to definitively categorise them as such. We also included LoF variants that were present in the gnomAD dataset but had not been reported in WD patients. This initial estimate predicted that approximately 1 in 2400 people would have pathogenic genotypes and would be at risk of developing WD, with 1 in 25 people being carriers of pathogenic variants.

This initial prevalence estimate did not take into account variant penetrance that may lead to people carrying WD genotypes either not expressing the disease or having milder phenotypes. We took steps to remove variants from our analyses that may be distorting prevalence estimates. These included *probable* low penetrant variants that based on few reports in the WD literature are at unexpectedly high frequencies, and *possible* low penetrant variants that were defined based on low VEST3 scores. After removal of these predicted low penetrant variants from our analysis the estimated prevalence of WD fell. Our rationale for removing the 13 variants classified as *probable* low penetrance is supported by a review of the WD literature. Given their frequencies in the gnomAD dataset the number of publications describing them in WD cohorts is much lower than expected (Abdelghaffar et al. 2008; Aggarwal et al. 2013; Cox et al. 2005; Davies et al. 2008; Garcia-Villarreal et al. 2000; Kim et al. 1998; Kroll et al. 2006; Lepori et al. 2007; Loudianos et al. 1999; Loudianos et al. 1998; Margarit et al. 2005; Mukherjee et al. 2014; Okada et al. 2000; Santhosh et al. 2006; Shah et al. 1997; Vrabelova et al. 2005). The publications reporting these variants also include data suggesting that some have low penetrance. The c.1947-4C>T variant is reported as a polymorphism in two publications (Kim et al. 1998; Okada et al. 2000) and appears to have been incorrectly classified as disease causing in the WDMD. The c.4021+3A>G (Santhosh et al. 2006) and p.Thr1434Met (Abdelghaffar et al. 2008) variants were identified in WD patients who were also homozygous or compound heterozygous for other *ATP7B* variants that could account for their phenotypes. A publication reporting p.Gly869Arg suggests that it has a more benign clinical course (Garcia-Villarreal et al. 2000), while p.Ile1230Val had an uncertain classification (Davies et al. 2008). Publications reporting the remainder of the *probable* low penetrant variants do not give clinical details of the patients involved, so that it is difficult to assess their pathogenicity. However, according to the ACMG standards and guidelines for the interpretation of sequence variants, “allele frequency greater than expected for disorder” is strong evidence for classification as a benign variant (Richards et al. 2015). Therefore, the gnomAD population data alone could be sufficient evidence to reclassify the variants we identified as being *probable* low penetrant to benign or likely benign. The functional characterisation of these variants would be useful for confirming that they are indeed low penetrant or non-causative. However, recent research suggests that current cell-based systems may not be accurate at measuring mild impairments in ATP7B function (Guttmann et al. 2018).

After removing variants with low VEST3 scores the predicted prevalence of WD genotypes fell further but because these variants were relatively infrequent the reduction was marginal. Although our analysis showed that the VEST3 algorithm performed well at discriminating between WD and non-WD *ATP7B* missense variants, no VEP algorithms are 100% accurate and hence the classification and removal of variants based purely on VEPs should be taken with caution. A recent report that analysed VEP algorithms suggests that these *in silico* analysis methods tend to over-estimate the pathogenicity of *ATP7B* variants unless thresholds are altered for the specific protein in question (Tang et al. 2019). When this is considered, there may be many more *ATP7B* variants that are incorrectly classified as disease causing and these may be distorting the predicted prevalence of the disease.

Based on our analysis of WD-*ATP7B* variant frequencies and considering the above strategies to account for low penetrant variants our final prediction for the population prevalence of WD is in the range of 1 in 17,000 to 1 in 20,000 of the global population with 1 in 65 to 1 in 70 as heterozygous carriers. It is of note that the predicted prevalence was not uniform across the 8 gnomAD subpopulations. The highest prevalence was observed in the Ashkenazi Jewish and East Asian subpopulations, both being close to 1 in 5000 with 1 in 36 heterozygous carriers. In the Ashkenazi Jewish population the most prevalent pathogenic variant was p.His1069Gln. This was also the most prevalent pathogenic variant in the European population and reflects the likely origin of this variant in the ancestors of Eastern Europeans (Gomes and Dedoussis 2016). In East Asians the most prevalent pathogenic variants were p.Thr935Met and p.Arg778Leu, both with similar allele frequencies.

Our prevalence estimate is higher than the traditional estimate of 1 in 30,000 but is not as high as other recent genetic estimates (Coffey et al. 2013; Collet et al. 2018; Gao et al. 2019). The recent Gao et al. study also used the gnomAD data to estimate the global prevalence of WD (Gao et al. 2019). However, while their method for estimating prevalence was similar, their analysis of penetrance was different to ours. Hence their final prevalence estimate of around 1 in 7000 is significantly higher (Gao et al. 2019). To address the issue of low penetrant variants, they used an equation reported by Whiffin et al (Whiffin et al. 2017) that calculates a maximum credible population allele frequency and filtered out all variants with allele frequencies higher than this. This method only removed 4 high frequency variants from their analysis. Recent studies from the UK and France also estimated relatively high carrier rates for WD using control populations (Coffey et al. 2013; Collet et al. 2018). Both studies filtered out some potentially low penetrant variants based largely on *in silico* computational analysis. However, unlike our study, none of these genetic studies used information from the WD literature to identify variants that are at too high a frequency in the population to be major contributors to WD genotypes. We have reanalysed the variant data from all 3 of the recent genetic studies using our criteria for classifying variants as *probable* or *possible* low penetrant. As the Gao et al. study used data from the same source as our study (Gao et al. 2019), reanalysis using our criteria returns a similar population prevalence of around 1 in 20,000. Many of the variants included in the Coffey et al. (Coffey et al. 2013) and Collet et al. (Collet et al. 2018) studies were classified as *probable* or *possible* low penetrant in our study and hence their removal results in significant reductions in the predicted population prevalence of WD, in the range of 1 in 47,000 in the UK and 1 in 30,000 to 54,000 in France (Supplementary Tables S5 and S6). These updated prevalence estimates are more closely aligned with what would be expected based on the clinical presentation of WD and suggest that reduced variant penetrance plays a much bigger role in the observed disparity between genetic and epidemiological studies (Gao et al. 2019). This parallels the conclusions of a recently published review on the epidemiology of WD (Sandahl and Ott 2019).

This study emphasises the difficulty in assigning WD prevalence from population datasets. Accurate prevalence estimates depend upon an assessment of the penetrance of individual genetic variants, not a straightforward task.

In conclusion, we have used NGS data to analyse the prevalence of WD in global populations, with a concerted approach to evaluating variant penetrance. This study highlights the importance of considering variant penetrance when assigning causality to genetic variants. Variants that have relatively high allele frequencies but low frequencies in patient cohorts are likely to have low penetrance. Large NGS datasets and improved VEP algorithms now allow us to evaluate with more accuracy the pathogenicity of genetic variants. The penetrance of *ATP7B* variants is likely to be on a spectrum: LoF variants are known to have high penetrance, whereas, some missense variants are thought to have lower penetrance (Chang and Hahn 2017). It would be valuable to determine the effects that low penetrant variants identified here have on ATP7B protein function and whether individuals carrying genotypes containing these variants have milder abnormalities of copper homeostasis, later onset or less severe forms of WD. Finally, this approach to predicting the prevalence of WD and penetrance of variants could be applied to other Mendelian inherited disorders.

## Acknowledgements

We would like to acknowledge Diane Wilson Cox and the curators of the Wilson Disease Mutation Database, University of Alberta for the use of data in this study.

## Supplementary Information

**Table S1. Disease causing variants identified in the Wilson Disease Mutation Database (WDMD).** Includes allele frequency data from the gnomAD population and subpopulations, number of references in the WDMD, whether referenced in Gomes and Dedoussis review (Ann Hum Biol (2016) 43:1-8) and VEST3 score.

**Table S2. Disease causing variants identified by a literature search between 2010 and 2017.** Includes allele frequency data from the gnomAD population and subpopulations, PubMed ID number, whether referenced in Gomes and Dedoussis review (Ann Hum Biol (2016) 43:1-8) and VEST3 score.

**Table S3.** *ATP7B* **loss of function variants and CNV deletions identified in gnomAD and ExAC databases.** Includes allele frequency data from the gnomAD population and subpopulations.

**Table S4. Combined WD-*ATP7B* variant allele frequencies, genotype frequencies and carrier rates in the gnomAD population** (not including additional non-WD reported LoF variants and CNVs).

**Table S5. Analysis of variants in UK control population: Coffey et al. Brain (2013) 136:1476-1487.**

**Table S6. Analysis of variants in French control population: Collet et al. BMC Medical Genetics (2018) 19:143.**

**Figure S1. Comparison of non-WD missense and WD missense *ATP7B* variants using 16 VEP algorithm scores.**

**Figure S2. Receiver operating characteristic (ROC) curve analysis was used to assess the effectiveness of 16 VEP algorithms to discriminate between WD missense and non-WD missense *ATP7B* variants**.

