## Supplementary for "*ATP7B* variant penetrance explains differences between genetic and clinical prevalence estimates for Wilson disease"

**Human Genetics**

Daniel F. Wallace*, James S. Dooley

* Corresponding author:

Institute of Health and Biomedical Innovation and School of Biomedical Sciences, Queensland University of Technology, Brisbane, Queensland, Australia.

 (DFW)

**Supplementary Material:**

Supplementary Table S4

Supplementary Results

Supplementary Figure S1

Supplementary Figure S2

Supplementary Reference

**Supplementary Table S4. Combined WD-*ATP7B* variant allele frequencies, genotype frequencies and carrier rates in the gnomAD population (not including additional non-WD reported LoF variants and CNVs)**

|  | **gnomAD** | | | | | | | | |
| --- | --- | --- | --- | --- | --- | --- | --- | --- | --- |
|  | **All** | **African** | **Ashkenazi Jewish** | **East Asian** | **European (non-Finnish)** | **European (Finnish)** | **Latino** | **South Asian** | **Other** |
| Pathogenic allele freq | 0.02004 | 0.01173 | 0.03005 | 0.02358 | 0.02278 | 0.01744 | 0.01629 | 0.01546 | 0.02142 |
| Pathogenic genotype freq | 0.00040 | 0.00014 | 0.00090 | 0.00056 | 0.00052 | 0.00030 | 0.00027 | 0.00024 | 0.00046 |
| Heterozygous genotype freq | 0.03927 | 0.02318 | 0.05830 | 0.04604 | 0.04452 | 0.03427 | 0.03204 | 0.03044 | 0.04191 |
| Pathogenic genotype carrier rate^†^ | 2491 | 7271 | 1107 | 1799 | 1927 | 3289 | 3770 | 4184 | 2180 |
| Heterozygous carrier rate^†^ | 25 | 43 | 17 | 22 | 22 | 29 | 31 | 33 | 24 |

^†^ Pathogenic genotype rate and heterozygous carrier rate are expressed as 1 in “n” of the population.

**Supplementary Results**

**Comparison of variant effect prediction (VEP) algorithms**

VEP algorithms are used extensively to predict whether amino acid substitutions (missense variants) are likely to alter protein function and hence contribute to disease. SIFT and Polyphen2 are two of the mostly widely used algorithms, however, in recent years newer algorithms have been developed. The output from wANNOVAR included results from 16 VEP algorithms. We tested the performance of these algorithms in discriminating between the 400 WD-*ATP7B* missense variants (identified in this study through literature review as associated with WD) and 786 missense variants (of 844 in total) that were identified in the gnomAD dataset but have not been previously reported in WD patients, termed non-WD-*ATP7B* missense variants. The scores for each of the algorithms were compared between the 2 groups (Supplementary Figure S1) and their performance in discriminating between the 2 groups assessed using ROC curve analyses (Supplementary Figure S2). Mean and median scores were compared between the two groups and the differences were statistically different for each algorithm (Supplementary Figure S1, t-test p<0.01, Mann Whitney test p<0.0001). ROC curve analyses revealed area under the ROC curves that ranged between 0.5399 and 0.8821 (Supplementary Figure S2).

The algorithm that performed the best at discriminating between WD and non-WD missense variants was VEST3 [1]. The median VEST3 score for WD missense variants was 0.957 compared with 0.404 for non-WD missense variants (Mann-Whitney test p<0.0001, AUROC 0.8821). None of the WD-*ATP7B* missense variants reported in the Gomes and Dedoussis review paper had VEST3 scores of less than 0.5 and only one variant with greater than 2 references in the WDMD had a VEST3 score of less than 0.5, indicating that the VEST3 score performs very well at discriminating between WD and non-WD *ATP7B* missense variants. We classified WD-*ATP7B* missense variants found in the WDMD and in our literature search as ‘*possible* low penetrance’ if they had a VEST3 score of <0.5 (Figure 1B). There were 11 such variants in the gnomAD dataset that were contributing to our initial estimates of WD prevalence (Table 3). Two of these variants were also classified as *probable* low penetrance in the previous analysis based on the number of publications.

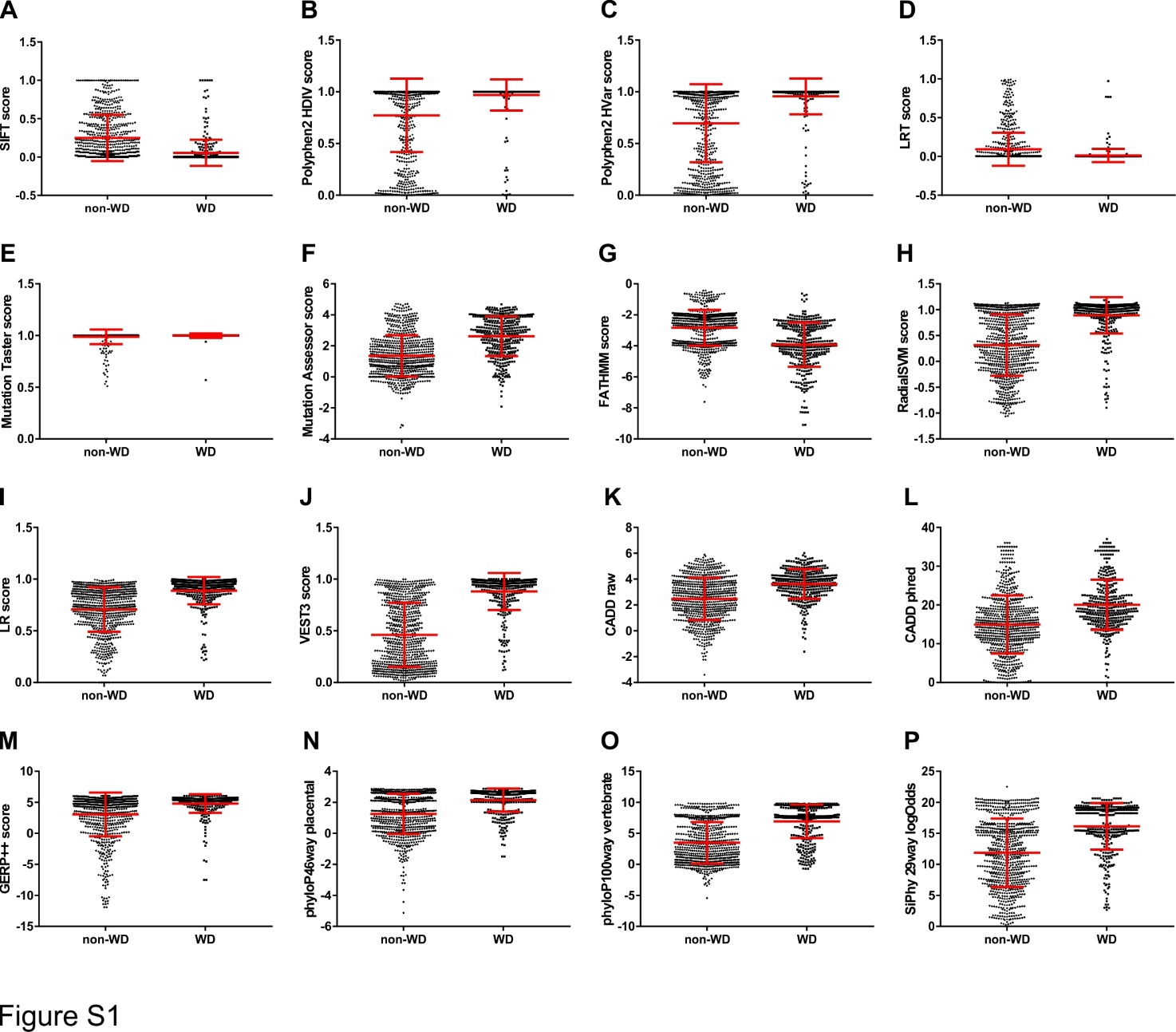

**Supplementary Figure S1.** Comparison of non-WD missense and WD missense *ATP7B* variants using 16 VEP algorithm scores. (A) SIFT, (B), Polyphen2 HDIV, (C) Polyphen2 HVAR, (D) LRT, (E) Mutation Taster, (F) Mutation Assessor, (G) FATHMM, (H) RadialSVM, (I) LR, (J) VEST3, (K), CADD raw, (L) CADD phred, (M) GERP++, (N) phyloP46way placental, (O) phyloP100way vertebrate, (P) SiPhy 29way logOdds. Data are presented as dot plots showing mean value and standard deviation.

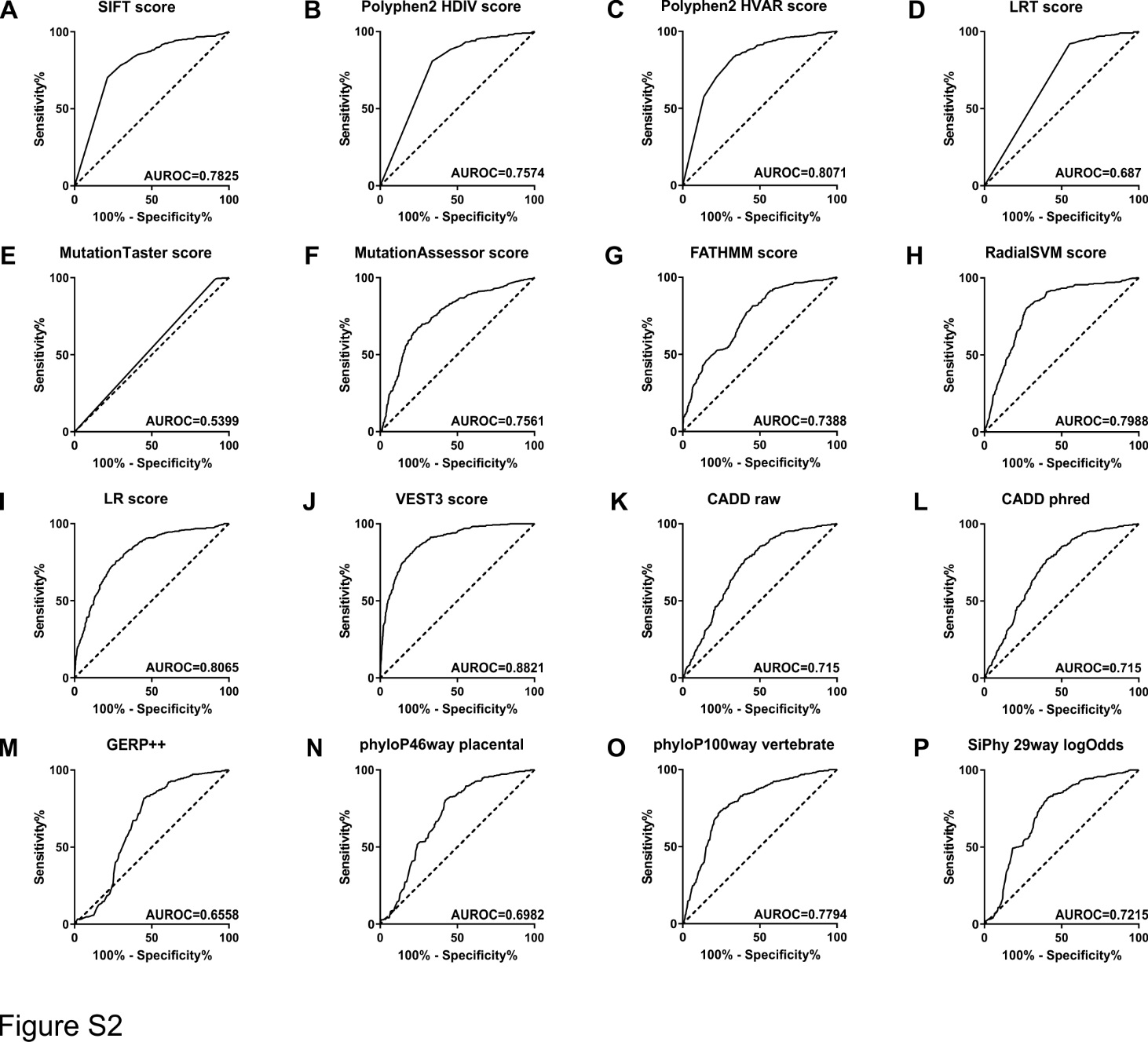

**Supplementary Figure S2.** Receiver operating characteristic (ROC) curve analysis was used to assess the effectiveness of 16 VEP algorithms to discriminate between WD missense and non-WD missense *ATP7B* variants. (A) SIFT, (B), Polyphen2 HDIV, (C) Polyphen2 HVAR, (D) LRT, (E) Mutation Taster, (F) Mutation Assessor, (G) FATHMM, (H) RadialSVM, (I) LR, (J) VEST3, (K), CADD raw, (L) CADD phred, (M) GERP++, (N) phyloP46way placental, (O) phyloP100way vertebrate, (P) SiPhy 29way logOdds. Area under the ROC curve (AUROC) is shown for each VEP algorithm.
